# Hyperbolic stratification of protein intrinsic disorder and structure-mediated interactions in the human protein interactome

**DOI:** 10.64898/2026.04.10.717685

**Authors:** Frank Hause, Oleksandr Sorokin, Stefan Hüttelmaier, Andrea Sinz

**Affiliations:** Department of Pharmaceutical Chemistry and Bioanalytics, Martin Luther University Halle-Wittenberg, Halle (Saale), Germany; Center for Structural Mass Spectrometry, Martin Luther University Halle-Wittenberg, Halle (Saale), Germany; Institute of Molecular Medicine, Section for Molecular Cell Biology, Faculty of Medicine, Martin Luther University Halle-Wittenberg, Halle (Saale), Germany

**Author notes:** Correspondence to: Frank Hause, Center for Structural Mass Spectrometry, Martin Luther University Halle-Wittenberg, Halle (Saale), Germany. Both authors contributed equally (Frank Hause, Oleksandr Sorokin).

## Abstract

Classical models of protein-protein interactions (PPIs) focus on stable, structure-driven interfaces between folded domains, yet recent work highlights the central role of intrinsic disorder and phase separation in shaping dynamic, multivalent associations. How these interaction modes are reflected in the large-scale organization of PPI networks remains unclear.

Here, we map the human interactome onto a hyperbolic representation, integrating sequence- and structure-derived features to test whether network organization reflects distinct molecular interaction strategies. Radial position defines a continuum: central proteins are enriched in folded domains, structural complexity, and post-translational modifications, whereas peripheral proteins show increased intrinsic disorder and liquid-liquid phase separation (LLPS) propensity. Angular organization further reveals communities structured by characteristic domain architectures or disorder-linked motifs.

Combined analysis of intrinsic disorder, LLPS propensity, and binding-mode diversity uncovers interaction patterns associated with distinct molecular functions and motif repertoires. Condensate-associated proteins span multiple communities while retaining shared short linear motif signatures. Together, these results show that the hyperbolic map links sequence composition, structural organization, and network topology, providing a framework to interpret protein interaction behavior and to guide functional analysis within the human interactome.

## Introduction

Biological systems consist of multiple, inter-dependent layers of molecular interaction networks that coordinate cellular function (1). Among these, protein-protein interaction (PPI) networks provide an important framework for understanding how cellular processes emerge from the complex interdependencies between proteins. A key feature of PPI networks is their hierarchical organization consisting of a small set of highly connected proteins forming integrated cores, whereas many proteins with lower connectivity occupy more peripheral positions (2, 3). Understanding how this network structure and its geometry relate to the molecular properties of the proteins embedded within it remains an important challenge in systems biology (4, 5).

Historically, the “protein folding dogma” held that a protein’s specific three-dimensional structure uniquely determines its function (6). Within this classical view, biological activity and interaction specificity were attributed to the formation of stable, complementary interfaces between folded domains (7). Consequently, PPIs were primarily conceived as rigid, well-defined binding events driven by the precise alignment of tertiary structural elements (8). This structure-centric paradigm successfully explained many enzymatic, structural, and signaling complexes governed by tight and specific domain-domain interactions.

The discovery of intrinsically disordered proteins (IDPs) and intrinsically disordered regions (IDRs) fundamentally challenged this dogma by revealing that structural order is not a prerequisite for function (9–12). IDPs engage in highly dynamic, context-dependent interactions that often involve short linear motifs (SLiMs) within flexible regions. These motifs mediate transient, multivalent, and low-affinity interactions that can be readily remodeled, providing a powerful mechanism for regulation and signaling. Importantly, subsets of IDPs and IDRs contain sequence features that promote liquid-liquid phase separation (LLPS), driving the selective recruitment of macromolecules into dense, dynamic condensates within the cellular environment (13, 14). Through such mechanisms, intrinsic disorder enables distributed and tunable connectivity within PPI networks, contrasting sharply with the discrete, lock-and-key logic of structured binding interfaces. Together, these discoveries expand the concept of protein interaction from static complexes to a continuum encompassing both stable, domain-mediated contacts and dynamic, disorder-driven assemblies.

Advances in network geometry have provided powerful approaches to study the organization of complex biological networks from a systems perspective. In particular, hyperbolic embeddings have emerged as an effective framework for representing complex networks (15, 16). In this approach, proteins are positioned on a curved ‘interaction landscape’, a geometric mapping defined by two coordinates: The radial coordinate describes the distance from the center, with more central proteins typically interacting with many partners, whereas peripheral proteins tend to have fewer interactions. The angular coordinate specifies the position along the map, analogous to the hands of a clock, and groups proteins involved in related biological processes or functions. Previous work has shown that, in the human PPI network, radial distance correlates with proteins’ evolutionary age, while angular proximity reflects functional similarity (17), indicating that the hyperbolic map captures underlying organizing principles that shape the large-scale structure of the proteome’s interaction network.

While hyperbolic maps capture evolutionary and functional organization, they have not yet been linked to the underlying molecular logic of protein interactions. In particular, it remains unknown whether the geometric organization of the PPI network reflects the structural and sequence-encoded mechanisms driving binding, such as the balance between folded and intrinsically disordered interaction modes. Clarifying this connection could reveal how network topology and molecular mechanisms co-evolve to shape the architecture of the human interactome (18).

Here, we address this question by integrating hyperbolic network geometry with sequence-derived features capturing disorder-mediated interaction mechanisms alongside structured protein elements and network topology. Using a hyperbolic map of the human PPI network, we test whether radial and angular organization stratifies proteins according to their molecular interaction strategies, with emphasis on intrinsic disorder, LLPS-associated sequence features, and SLiMs. By linking network geometric organization to sequence architecture, structural composition, and functional context, this study provides a unified framework for how molecular interaction mechanisms are encoded in the large-scale structure of the human interactome.

## Results

### Hyperbolic PPI network organization and structural protein characteristics

To investigate how molecular interaction mechanisms are reflected in the geometric organization of the human PPI network, we first constructed a hyperbolic representation of the human PPI network consisting of 200,766 high-confidence interactions between 11,693 proteins. In this representation, the radial coordinate reflects how central a protein is on the map, with smaller radial values indicating proteins that interact with many partners. Consistent with this interpretation, radial position showed a strong inverse association with node degree (Spearman’s ρ = −0.981, p << 10^-300^) (Supplementary Figure S2a). In contrast, radial position was strongly positively associated with local fractal dimension (LFD; a measure of the complexity of a protein’s local interaction neighborhood), which increases from central to more peripheral regions of the network (ρ = 0.931, p << 10^-300^) (Supplementary Figure S2b).

Network communities, defined as groups of proteins that are more densely connected to each other than to the rest of the network, were identified using a random walk (RW) approach. In this framework, a hypothetical walker that moves a limited number of steps along the network is more likely to remain within densely connected regions, which are consequently identified as communities. Pairwise angular distances were significantly smaller within RW communities than between communities, with within-community pairs strongly concentrated near zero and exhibiting a pronounced tail, whereas between-community pairs were distributed more evenly across the full angular range (Wilcoxon p << 10^-300^) (Supplementary Figure S2c), indicating that community structure is reflected as tight clustering along the angular dimension of the hyperbolic map. Likewise, true interaction edges occurred at significantly smaller hyperbolic distances than degree-matched non-interacting pairs (Wilcoxon p << 10^-300^) (Supplementary Figure S2d), showing that proximity on the hyperbolic map is strongly associated with interaction likelihood.

Radial organization of the human PPI network is associated with structured protein architecture after controlling for degree. Although the correlation between protein length and hyperbolic radius remained significant, it was minor (partial Spearman’s ρ = −0.021, p = 0.026) (Fig. 1a). In contrast, smaller radial positions were significantly associated with a higher number of post-translational modifications (PTMs) per protein when controlling for degree (part. Spearman’s ρ = −0.192, p = 7.08 × 10^-95^) (Fig. 1b). Similar trends were observed for domain number (ρ = −0.111, p = 1.50 × 10^-32^) (Fig. 1c) and the overall number of annotated structural elements (ρ = −0.0614, p = 5.66 × 10^-11^) (Fig. 1d).

**Figure 1.**
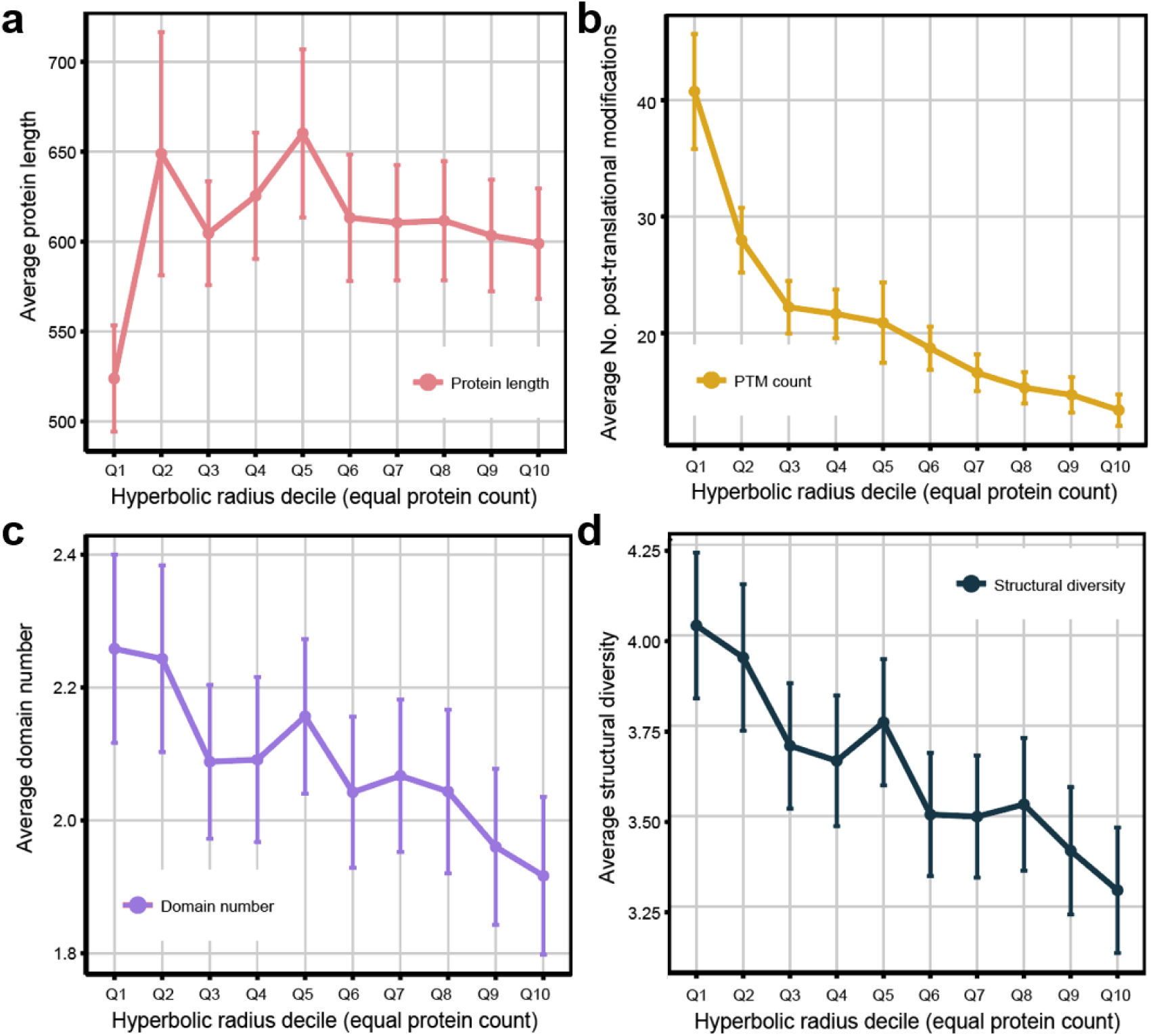
Radial position stratifies protein architectural complexity in the hyperbolic projection of the human PPI network. (a) Average protein length shows only modest variation across hyperbolic radius deciles, with no consistent monotonic trend. (b) The average number of PTMs per protein decreases markedly from central to peripheral regions. (c) The average number of annotated domains per protein shows a gradual decline with increasing radial distance. (d) Structural diversity, defined as the number of distinct annotated structural elements per protein, is highest in the network core and progressively decreases toward the periphery. All values are shown as mean (± standard error of mean [s.e.m.]) across proteins within each radial decile (Q1-Q10, equal protein counts).

Protein domain diversity differs strongly across RW communities (Kruskal-Wallis χ² = 868.97, df = 44, p = 1.04 × 10^-153^), indicating that communities partition proteins into structurally distinct groups. Community-level enrichment analysis revealed highly specific concentration of characteristic InterPro families within individual communities (Supplementary Table S1). For example, community 8 is strongly enriched for rhodopsin-like seven-transmembrane GPCR (G-protein coupled receptor) domains (OR = 159.17, p = 5.86 × 10^-188^), community 2 for immunoglobulin-like architectures (OR = 13.65, p = 1.03 × 10^-135^) and SH2 domain-containing proteins (OR = 48.41, p = 2.23 × 10^-54^), and community 4 for histone-fold domains (OR = 43.62, p = 3.01 × 10^-65^). Many of these enrichments are extremely strong, showing that several communities correspond to sharply delimited structural protein classes rather than arbitrary graph partitions.

### Organization of intrinsic disorder in radial and angular dimension

We next investigated whether intrinsic disorder is reflected in the organization of the human PPI network. Disorder levels were heterogeneously distributed, with most proteins exhibiting low disorder fractions and a minority displaying high disorder (Fig. 2a). AIUPred (19) systematically predicted higher IDR fractions than AlphaFold2 (20, 21), with overall concordance between methods (Spearman’s ρ = 0.589, p << 10^-300^, R² = 0.312) but increasing divergence at higher disorder levels.

**Figure 2.**
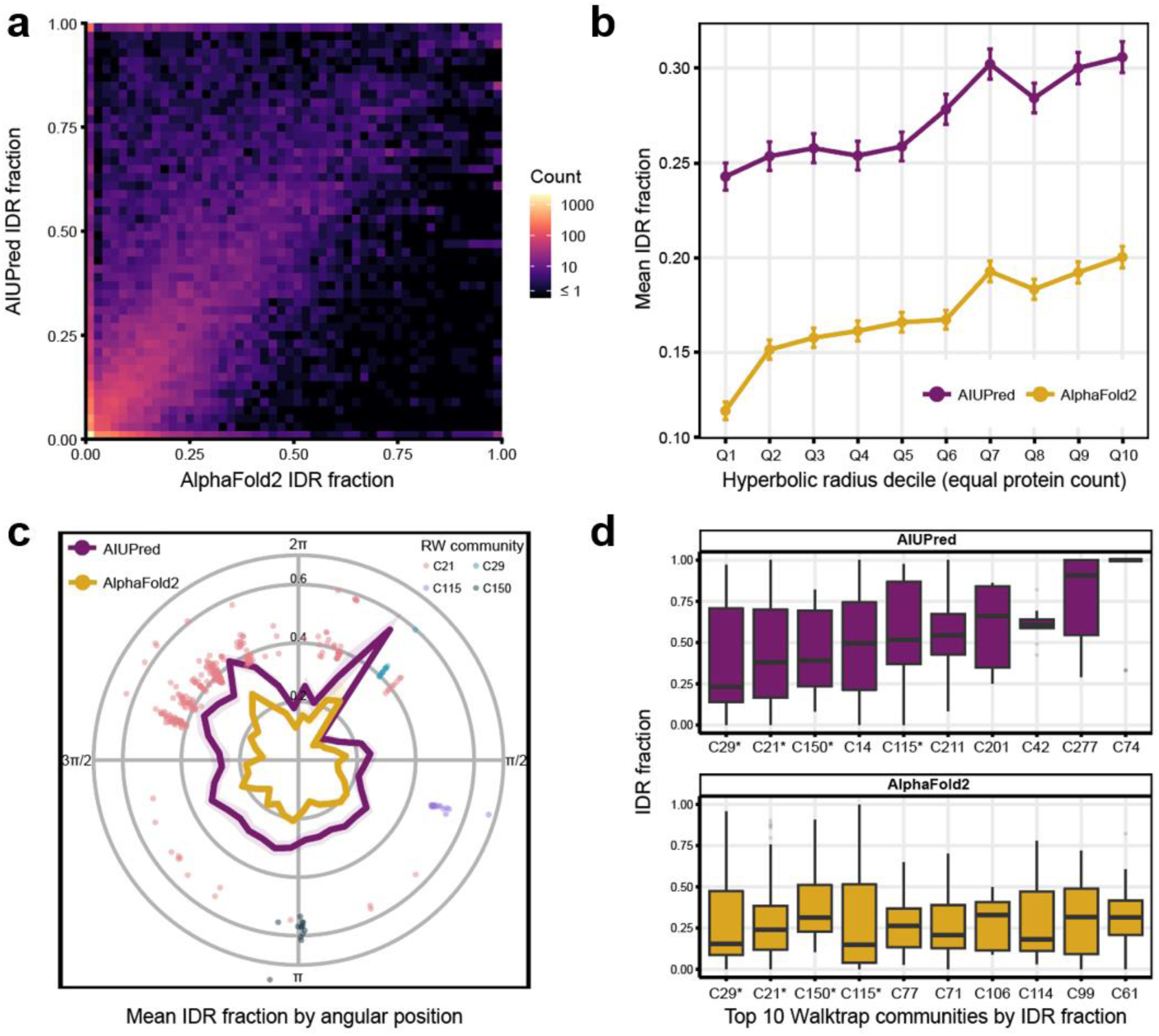
Intrinsic disorder is radially and angularly organized in the hyperbolic human PPI network. (a) Density of IDR fraction per protein estimated by AIUPred and AlphaFold2 across all proteins in the human PPI network. Values showed strong concordance at low disorder fractions, while at higher IDR levels the agreement gradually decreases. (b) Mean IDR fraction across hyperbolic radius deciles (Q1-Q10, equal protein counts). Both AIUPred (purple) and AlphaFold2 (gold) show a consistent increase in disorder toward peripheral regions. Values are shown as mean ± s.e.m. (c) Angular distribution of mean disorder fraction for AIUPred (purple) and AlphaFold2 (gold). Intrinsic disorder is unevenly distributed across angular sectors, with distinct peaks indicating preferential localization of IDR-rich proteins. Selected IDR-rich RW communities, identified by both AIUPred and AlphaFold2 among those with the highest IDR fraction, are shown at their corresponding angular positions. (d) Distribution of IDR fractions across RW communities with the highest predicted intrinsic disorder. Boxplots for AIUPred (purple) and AlphaFold2 (gold) highlight substantial heterogeneity between communities and prediction methods.

Projection onto the hyperbolic map revealed a non-random radial organization of protein IDR fraction. When controlled for degree, mean IDR fraction increased with radial distance from the network center (AIUPred: Spearman’s ρ = 0.067, p = 3.9 × 10^-13^; Alphafold2: Spearman’s ρ = 0.043, p = 3.01 × 10^-6^) (Fig. 2b), indicating preferential localization of IDPs toward the periphery. Additionally, the IDR fraction of proteins was also structured along the angular dimension. Sector-specific enrichments were observed, with localized peaks in disorder fraction defining discrete angular regions (Fig. 2c).

At the RW community level, IDR fraction differed markedly (Fig. 2d). Among the ten RW communities with the highest predicted IDR fractions, four were shared between AIUPred and AlphaFold2-based predictions. Functional over-representation analysis (ORA) of these shared communities revealed a broad functional spectrum of IDR-rich protein groups, including mRNA binding (C21: GO:0003729, adj. p = 2,35 × 10^-86^), skin development (C29: GO:0043588, adj. p = 8.21 × 10^-14^), maintenance of gastrointestinal epithelium (C115: GO:0030277, adj. p = 3.04 × 10^-6^), to exocytosis (C150: GO:0006887, adj. p = 7.51 × 10^-9^). Notably, enrichment terms in IDR-rich RW communities were predominantly associated with interface-related processes as captured in transport- or binding-related gene sets (Supplementary Table S2).

### Integration of LLPS propensity and binding mode multiplicity in hyperbolic space

To assess whether phase-separation-associated features follow the geometric organization of intrinsic disorder on the hyperbolic map, we quantified LLPS-related sequence properties using FuzDrop (22–24). This includes the probability of disorder-to-order binding (pDO), reflecting the likelihood that a disordered region adopts a folded structure upon interaction, and disorder-to-disorder binding (pDD), indicating retention of conformational flexibility in the bound state. In addition, a global LLPS propensity score, p(LLPS), estimates the likelihood of a protein to undergo LLPS. We further considered the multiplicity of binding modes (MBM), which captures the likelihood of a protein’s residues to switch between different interaction modes depending on its partner or cellular context. For example, regions with high MBM may remain disordered in transient interactions but adopt structured conformations under specific conditions, and in some cases are prone to aggregation.

Global LLPS propensity (part. Spearman’s ρ = 0.064, p = 5.55 × 10^-12^), pDD fraction (part. Spearman’s ρ = 0.083, p = 6.58 × 10^-19^), and pDO fraction (part. Spearman’s ρ = 0.078, p = 3.19 × 10^-17^) increased towards the periphery of the hyperbolic map (Fig. 3a), mirroring the radial increase of mean IDR fraction. In contrast, MBM fraction showed only weak radial variation (part. Spearman’s ρ = 0.04, p = 1.28 × 10^-5^), indicating that interaction versatility is not strongly coupled to disorder content.

**Figure 3.**
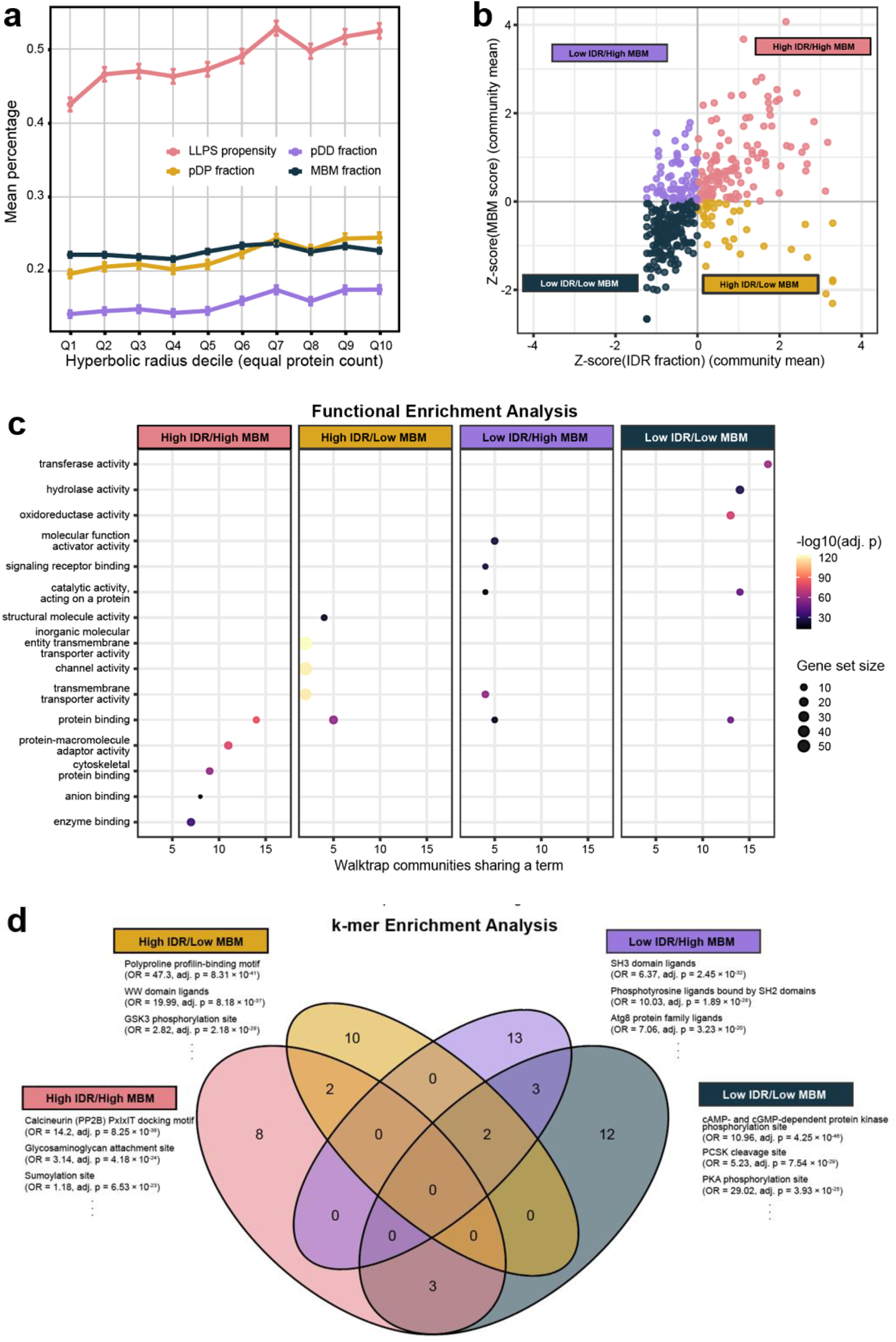
Interaction regimes emerge from intrinsic disorder and binding mode multiplicity in the human interactome. **(a)** Mean LLPS propensity (red), pDD fraction (purple), pDP fraction (gold), and MBM fraction (dark blue) across hyperbolic radius deciles (Q1-Q10, equal protein counts). LLPS propensity and disorder-related fractions increase toward the periphery, whereas MBM fraction shows only weak radial variation. Values are shown as mean ± s.e.m. **(b)** RW communities positioned by mean IDR fraction (x-axis, Z-score) and mean MBM fraction (y-axis, Z-score). Communities span all four quadrants defined by high or low disorder content and high or low MBM content, indicating that these features represent largely independent axes of interaction behavior. **(c)** Functional enrichment across the four interaction groups defined in (b). Each panel summarizes over-represented Gene Ontology molecular function terms across communities within a group. Distinct functional profiles separate macromolecule binding, enzymatic activity, signaling, and transport-related processes. **(d)** *k*-mer enrichment across the four interaction groups. Venn diagram showing overlap of annotated SLiMs between groups defined in (b). Each group displays a distinct motif repertoire consistent with specific interaction strategies.

At the RW community level, mean IDR and MBM fraction defined two largely independent axes. RW communities were distributed across four groups of high or low IDR and MBM fraction (defined relative to the global mean, hereafter termed interaction groups) (Fig. 3b), suggesting distinct interaction regimes within the human PPI network. Functional enrichment analysis showed clear separation of molecular roles (Fig. 3c). High IDR/high MBM communities (n = 118) were enriched for macromolecule binding and protein-protein interaction functions, consistent with multivalent interaction hubs. High IDR/low MBM communities (n = 36) were mainly associated with transmembrane functions. Low IDR/high MBM communities (n = 68) were enriched for signaling and adaptor functions, whereas low disorder/low MBM communities (n = 166) were linked to specialized enzymatic activities and catalytic processes.

To determine whether the interaction groups are enriched for specific sequence features that may drive interactions along the IDR–MBM axis, we performed *k*-mer enrichment analysis, identifying overrepresented short amino acid sequence patterns. Mapping these enriched *k*-mers to annotated short linear motifs (SLiMs) revealed distinct motif repertoires for each interaction group, with only minor overlap (Fig. 3d; Supplementary Table S3).

### Condensate-associated communities and their sequence signatures in the human protein interactome

Given that LLPS-associated sequence features were enriched toward peripheral regions of the hyperbolic map and defined distinct interaction patterns, we next investigated whether these properties are reflected at the level of biomolecular condensates, which are dynamic membraneless compartments formed by phase separation. To this end, we tested whether RW communities are enriched for proteins annotated to specific biomolecular condensates.

Across all RW communities, we identified 91 significant community-condensate enrichments after multiple testing correction (Supplementary Table S3), indicating that condensate-associated proteins are not randomly scattered on the hyperbolic map. Among the most prominently enriched condensates were the nucleolus (significantly enriched among n = 9 communities), stress granules (n = 7 communities), P-bodies (n = 5 communities), and nuclear speckles (n = 3 communities) (Fig. 4a), suggesting that condensate organization is captured as a distributed, yet structured feature of network topology.

**Figure 4.**
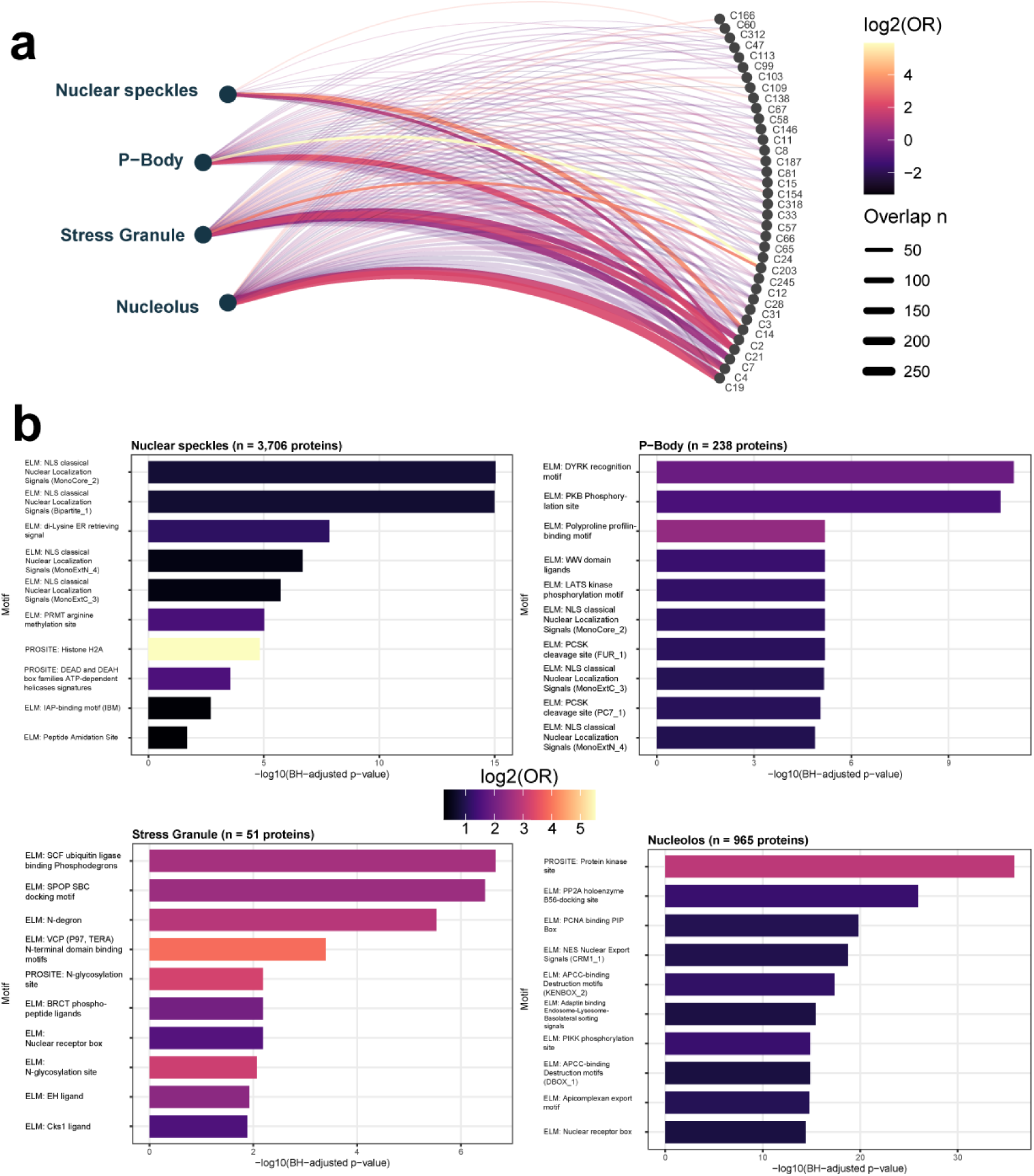
Biomolecular condensates map to distributed network communities and exhibit distinct sequence motif signatures. (a) Enrichment of biomolecular condensates across RW communities. Edges connect condensates (left) to significantly enriched communities (right), with edge color indicating enrichment (log2 odds ratios [OR]) and edge width representing overlap size (number of shared proteins). Condensate-associated proteins are distributed across multiple RW communities rather than confined to single modules, indicating a structured but distributed organization within the interactome. (b) Motif overrepresentation analysis of proteins associated with selected condensates. Bar plots show significantly enriched motifs annotated in ELM (23, 24) and PROSITE (25) for nuclear speckles, P-bodies, stress granules, and nucleoli, with significance represented as – log10(adj. p value) and color indicating effect size (log2 OR). Distinct motif repertoires are observed across condensates, including enrichment of nuclear localization signals in nuclear speckles and nucleoli, phosphorylation-related motifs in P-bodies, and degron-associated motifs in stress granules.

We next asked whether proteins associated with these condensates exhibit characteristic SLiM signatures. Motif overrepresentation analysis revealed distinct motif repertoires across condensates (Fig. 4b; Supplementary Table S4). Nuclear speckle and nucleolar proteins were strongly enriched for nuclear localization signals, consistent with their subcellular localization. In contrast, stress granule-associated proteins showed enrichment for degron-related motifs, indicating a potential link between dynamic turnover and condensate-associated interaction networks. P-body-associated proteins displayed enrichment for phosphorylation-related motifs, reflecting regulatory control via signaling pathways.

Finally, we examined whether these condensate-associated motif signatures relate to the sequence-level interaction regimes identified by *k*-mer enrichment. While the overlap was not uniform, consistent tendencies were observed (Supplementary Fig. S3). Nuclear speckles showed an enrichment of *k*-mers associated with higher IDR fraction, whereas P-bodies and nucleoli tended to be associated with *k*-mers reflecting comparatively lower disorder content. Stress granules were relatively depleted of *k*-mers linked to low IDR and low MBM fraction.

## Discussion

Hyperbolic network geometry provides a powerful framework to relate sequence-derived features to the large-scale organization of the human PPI network. Across multiple levels of analysis, position on the hyperbolic map and community structure capture distinct yet complementary aspects of protein interaction strategies. Importantly, this representation provides a useful framework for interpreting protein function within a network context. By positioning proteins along a continuum from central, structured interaction hubs to peripheral, disorder-driven interaction specialists, the hyperbolic map highlights proteins with distinct interaction behaviors. For experimental studies, hyperbolic PPI network maps may help to prioritize candidates based on inferred interaction mode, provide context for functional interpretation of less well-characterized proteins, and point to regions of the interactome that are likely influenced by dynamic, context-dependent interactions such as LLPS.

At the global level, the radial position reflects a gradient in protein architecture and evolutionary age. Proteins located toward the hyperbolic core exhibit increased mean domain content, higher structural diversity, and elevated numbers of post-translational modifications (Fig. 1). This stratification is consistent with the seminal work by Alanis-Lobato et al. (17) showing that the hyperbolic radius of a PPI embedding approximates protein evolutionary age, with older, more conserved proteins occupying central positions of the hyperbolic map and younger proteins residing at the periphery. The enrichment of multi-domain architectures and regulatory modification sites in the core is therefore consistent with an evolutionary scenario of domain accretion and functional refinement (25), in which duplication, recombination, and fusion of domains progressively expand structural and functional complexity.

Whereas structured protein architecture predominates in central regions of the hyperbolic map, peripheral proteins are characterized by increased intrinsic disorder (Fig. 2b). Importantly, intrinsic disorder reflects not the absence of annotated domains, but the ability of proteins to sample conformational ensembles that enable context-dependent and multivalent interactions. This structural plasticity provides a mechanistic basis for the interaction versatility observed in IDPs (26). Consistent with this, LLPS-associated features closely track IDR content and inversely relate to the abundance of structured elements, supporting a model in which disorder-driven interaction modes operate as a complementary organizational principle to domain-mediated interactions within the human interactome. Notably, disorder-driven and transient interactions are typically underrepresented in large-scale interaction datasets such as high-confidence STRING networks, which prioritize stable and recurrent contacts, suggesting that the observed organization likely represents a conservative estimate of the contribution of dynamic interaction modes.

At the level of interaction regimes, the integration of intrinsic disorder and MBM (22–24) further refines this picture. Whereas disorder-related features determining LLPS propensity follow the radial gradient, MBM varies more independently (Fig. 3a), pointing to additional layers of regulation that fine-tune interaction behavior beyond intrinsic disorder alone. This decoupling suggests that interaction promiscuity in IDPs is not solely determined by conformational flexibility, but is further modulated by local sequence patterning, post-translational regulation, and the repertoire of accessible binding partners.

At the community level, this interplay stratifies molecular function along the IDR–MBM axis. Proteins with low intrinsic disorder and low binding mode multiplicity are predominantly associated with enzymatic and catalytic functions, whereas increasing disorder and MBM are linked to signaling, regulatory, scaffold, and interface-related processes (Fig. 3c). This functional separation reflects fundamental constraints on molecular mechanisms. Enzymatic proteins typically rely on highly ordered architectures in which precise domain and subdomain arrangements are required to form catalytically competent active sites. Even subtle perturbations in structural alignment can disrupt activity, underscoring the necessity of rigid and well-defined conformational states (27). In contrast, regulatory proteins, such as transcription factors, exhibit a markedly higher tolerance toward structural flexibility. Perturbed or disordered activation domains can maintain functionality despite sequence rearrangements or mutations, as long as key physicochemical features are preserved (28, 29). This tolerance enables adaptive and context-dependent interactions, allowing the same protein to engage multiple partners across different cellular states. Consistent with this, the IDR–MBM stratification suggests that structural rigidity becomes progressively less critical for binding- and interface-dominated functions, where functional specificity is achieved through flexible, multivalent, and context-dependent interaction modes rather than fixed structural conformations.

Biomolecular condensates add a further layer of organization to this picture. We show that condensate-associated proteins exhibit distinct SLiM signatures that partially align with the sequence-derived interaction regimes identified by *k*-mer analysis of IDR–MBM-stratified protein communities (Fig. 4b). This correspondence indicates that condensate formation is not driven by isolated sequence features, but reflects broader interaction principles embedded in protein sequence architecture. A mechanistic explanation is provided by the stickers-and-spacers framework (30). Condensates arise from the spatial arrangement of interaction-promoting motifs (“stickers”), such as SLiMs and charged residues, embedded within intrinsically disordered “spacers”, enabling tunable multivalency and interaction strength. Within this framework, our results support a model in which compatible interaction features distributed across the protein interactome dynamically converge to form condensates, rather than being restricted to a small number of pre-defined network modules. Notably, condensate-associated proteins are distributed across multiple RW communities rather than confined to single modules (Fig. 4a), suggesting that condensate formation integrates interaction features across topological boundaries. This distributed organization may in part reflect the use of a high-confidence interactome, which prioritizes stable interactions while underrepresenting transient and low-affinity contacts that are central to condensate formation and IDR-mediated binding. Consequently, dynamic interaction layers are likely only partially captured in the current network, further supporting the view that condensates emerge from interaction regimes that extend beyond static PPI architectures. Consistent with this dynamic view, multivalent condensates form interaction networks in which a majority of proteins remain mobile while a smaller fraction acts as transient cross-links, giving rise to emergent mechanical behavior governed by continuous binding and unbinding dynamics (31). As such, condensate-associated interaction modes likely extend beyond the static representations captured in curated PPI networks and are further shaped by proteoform diversity, post-translational regulation, and spatial compartmentalization.

Beyond a simple order-disorder axis, the hyperbolic map reveals that protein interactions are shaped by a diverse spectrum of structural and sequence-encoded features, including SLiMs and multivalent interaction motifs, which give rise to dynamic molecular assemblies such as biomolecular condensates. These findings indicate that the human PPI network is realized by a continuum of interaction strategies that integrate stable, domain-mediated architectures with flexible, disorder-driven mechanisms essential for cellular function. By linking geometric network organization to molecular interaction principles, this study provides a conceptual basis for understanding how these diverse strategies are coordinated within the protein interactome and how they may be rewired across cellular states and disease contexts.

## Materials and Methods

### Human PPI network and hyperbolic embedding

The human PPI network analyzed in this study was constructed from the STRING database (v12.0) (32, 33). Only high-confidence interactions with a combined score greater than 0.900 were retained. Proteins were represented as nodes and interactions as undirected, unweighted edges. Self-interactions and duplicated edges were removed, and the analysis was restricted to the largest connected component (LCC) of the resulting network.

The network was embedded into two-dimensional hyperbolic space using the Mercator algorithm (34). Mercator implements a statistical inference framework based on the Popularity-Similarity Optimization (PSO) model, which assumes that network connectivity arises from a trade-off between node popularity and similarity in hyperbolic space. For each protein, Mercator (34) estimates the two hyperbolic coordinates, namely radial and angular. Based on these coordinates, pairwise hyperbolic distances and angular differences between all proteins were calculated to quantify geometric proximity within the embedded human PPI network.

### Network topology analysis

To characterize the structural organization of the human PPI network and to validate the hyperbolic embedding, vertex degree was computed using the igraph package (35). In addition, the local fractal dimension was calculated to quantify the increase in network complexity around individual nodes (36). Consistent with hyperbolic network geometry, LFD is expected to peak in peripheral proteins and exhibit a negative correlation with vertex degree. RW community structure was inferred using the Walktrap community detection algorithm (37) as implemented in the igraph package (35). The random-walk length parameter was systematically evaluated across a range of step sizes (from 2 to 50), and a step length of 18 was selected based on maximal modularity observed in the modularity versus step-size profile yielding 390 communities (Supplementary Figure S1).

### Community-wise functional over-representation analysis

ORA was performed independently for each RW-defined community using Gene Ontology (GO:BP, GO:MF, and GO:CC) (38, 39) (release Jan 23, 2026) and Reactome (40) (v95, release Dec 3, 2025) gene set annotations. The background universe was defined as all Entrez-mapped genes present in the LCC of the human STRING PPI network after preprocessing. Enrichment significance was assessed using a hypergeometric test with Benjamini-Hochberg (41) correction for multiple testing. Gene sets were restricted to sizes between 10 and 500 genes to exclude overly specific and overly broad annotations.

### Protein annotation from InterPro

Protein annotations were obtained for all STRING proteins within the LCC by mapping ENSP identifiers to Ensembl gene IDs and UniProt (42) Swiss-Prot accessions using biomaRt (43). Protein domain architecture was assessed via the InterPro (44, 45) REST API based on the mapped UniProt accessions. We quantified strict domain number per protein and broader architectural complexity as the number of unique entries of structural elements.

### Intrinsic disorder estimation using AIUPred and AlphaFold2

Intrinsic disorder was estimated for all proteins contained in the human STRING LCC using two independent residue-level resources, AIUPred (19) and AlphaFold2 predicted protein structures obtained from AlphaFold Protein Structure Database v6 (20, 21). In AIUPred (19), residues with disorder scores > 0.5 were classified as disordered. In AlphaFold2 (20, 21), disorder was approximated from structural confidence, defining residues with pLDDT values < 50 as disordered. Contiguous IDRs were defined as stretches of at least 10 consecutive disordered amino acid residues (46).

### LLPS propensity profiling using FuzDrop

LLPS-associated sequence features were quantified for all proteins in the human STRING LCC using residue-resolved predictions from FuzDrop (22–24). For each protein, disorder-to-order probability (pDO), disorder-to-disorder probability (pDD), multiplicity of binding modes (MBM), and the global LLPS score, p(LLPS), were extracted. Residues exceeding defined thresholds (pDO > 0.6, pDD > 0.6, MBM > 0.65) were classified as high-propensity. Consecutive stretches of at least 10 such residues were considered phase-separation-relevant segments.

### SLiM annotation from ELM and PROSITE

SLiMs were annotated using the Eukaryotic Linear Motif (ELM) resource (47, 48). UniProt accessions derived from the STRING node mapping were used to query the ELM server, which provides motif annotations in tabular format for individual proteins. For each protein, motif annotations were summarized by counting the total number of motif matches as well as the number of unique motif classes, motif instances, and motif accessions.

Additional motif annotations were retrieved from the PROSITE database (49) to complement SLiM coverage.

### Biomolecular condensate annotation from CD-CODE

Biomolecular condensate annotations were retrieved from the Crowdsourcing Condensate Database and Encyclopedia (CD-CODE) database (v2.2, release Mar 18, 2026) (50), which provides curated assignments of proteins to biomolecular condensates based on literature evidence and computational integration.

### *k*-mer enrichment

To characterize sequence-level determinants associated with distinct interaction regimes, *k*-mer enrichment analysis was performed on protein sequences grouped by Z-score transformed IDR and MBM fraction. For each group, protein sequences were compiled and analyzed separately across four sequence stretch properties (high IDR + high MBM; high IDR + low MBM; low IDR + high MBM; low IDR + low MBM), where high and low denote values above and below the global mean, respectively. All possible contiguous *k*-mers of length *k* = [3, …, 12] were extracted from each sequence using a sliding-window approach. For each group and each value of *k*, *k*-mer frequencies were quantified across all sequences, and the 100 most frequent *k*-mers were retained for downstream analysis.

### Motif overrepresentation analysis

To identify sequence motifs associated with distinct interaction regimes, overrepresentation analysis was performed on enriched *k*-mers using motif definitions from ELM (47, 48) and PROSITE (49) represented as regular expressions.

For each interaction group, the set of enriched *k*-mers was used as the foreground. A background universe was constructed from all unique *k*-mers extracted from the corresponding group-specific protein sequences across the same k-range, excluding foreground *k*-mers.

Motif occurrences were quantified by matching each motif regular expression against the foreground and background *k*-mer sets. For each motif, enrichment was assessed using one-sided Fisher’s exact test based on counts of matching and non-matching *k*-mers in foreground and background sets. Odds ratios (OR) were computed to estimate effect sizes.

## Statistical Analysis

All summary statistics are reported as mean ± s.e.m., unless stated otherwise. Statistical significance was assessed using appropriate tests as described for each analysis. Where multiple comparisons were performed, p values were adjusted using the Benjamini-Hochberg procedure (41). Adjusted p < 0.05 was considered statistically significant.

## Acknowledgements

This work was supported by the Deutsche Forschungsgemeinschaft (DFG) (RTG 2467, project number 391498659 “Intrinsically Disordered Proteins-Molecular Principles, Cellular Functions, and Diseases”, CRC 1664, project number 514901783 “SNP2Prot - Plant Proteoform Diversity”, and CRC 1423, project number 421152132 “Structural Dynamics of GPCR Activation and Signaling”). AS and SH acknowledge additional financial support by the DFG (RTG 2751 “InCuPanC”, project number 449501615; RU 5433 “RNA in Focus” project number 468534282, INST 271/404-1 FUGG; INST 271/405-1 FUGG; INST271/528-1 FUGG). Additionally, AS acknowledges financial support by the Federal Ministry for Economic Affairs and Energy (BMWi, ZIM project KK5096401SK0), the European Regional Development Funds for Saxony-Anhalt (grant numbers EFRE ZS/2024/01/183756 and ZS/2024/01/183619), and the Martin Luther University Halle-Wittenberg (Center for Structural Mass Spectrometry).

## Author Contributions

F.H. conceived the study. F.H. and O.S. designed the study, performed data acquisition, analysis, and visualization, and wrote the manuscript (first draft and revision). S.H. and A.S. contributed to data analysis and critically reviewed the manuscript. S.H. and A.S. supervised the study and acquired funding.

## Data Availability Statement

All data used in this study were obtained from publicly available tools and databases, including AIUPred (19), AlphaFold2 (20, 21), FuzDrop (22–24), STRING (32, 33), Gene Ontology (38, 39), Reactome (40), UniProt (42), InterPro (44, 45), ELM (47, 48), PROSITE (49), and CD-CODE (50), as detailed in the Materials and Methods. The R/Python code for data retrieval and processing is deposited at https://github.com/DataScienceFH/HyperbolIDRInteractome.

