## Supplementary figures and images for "Hyperbolic stratification of protein intrinsic disorder and structure-mediated interactions in the human protein interactome"

### Supplementary Material

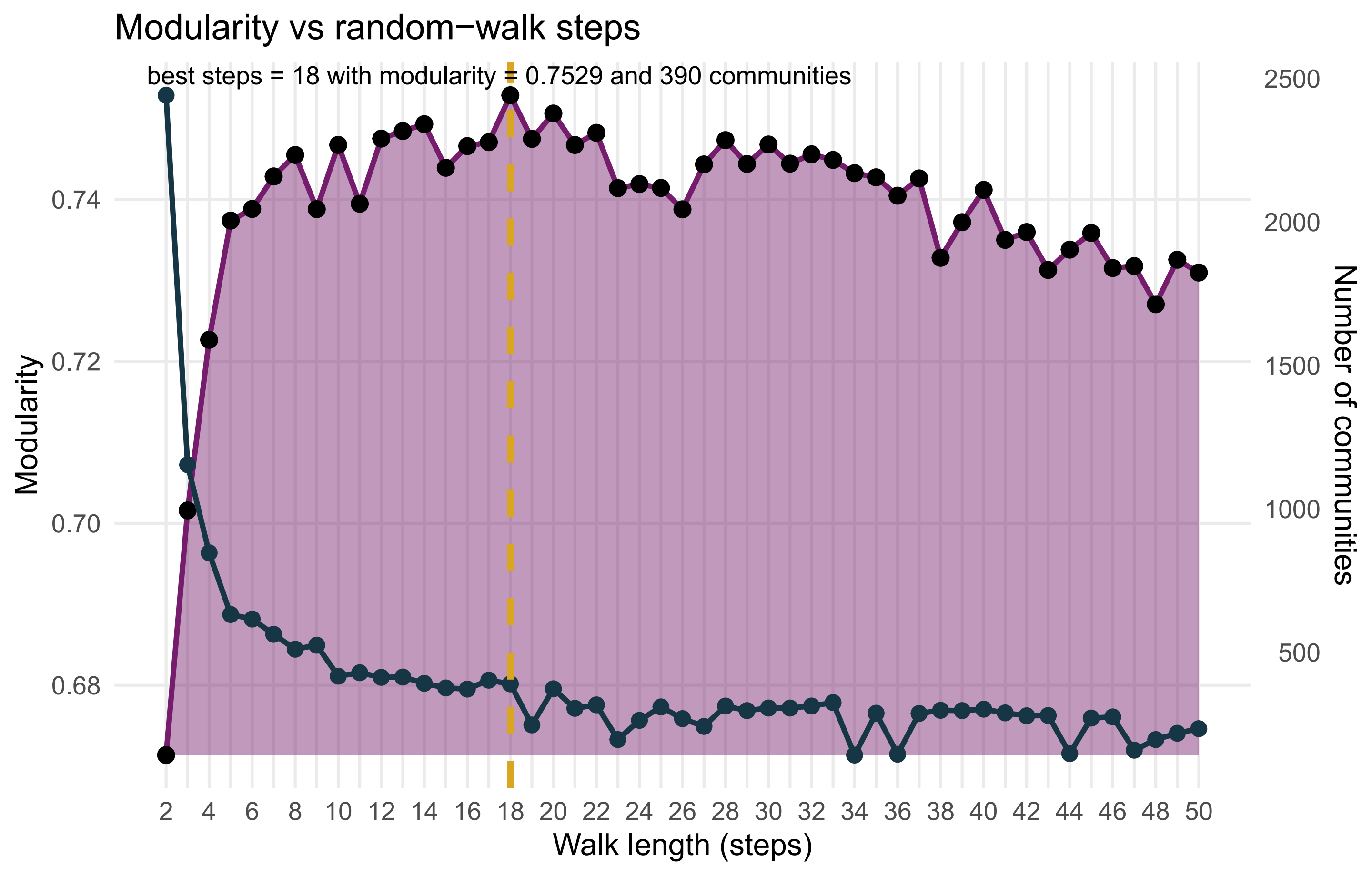

### Supplementary Material

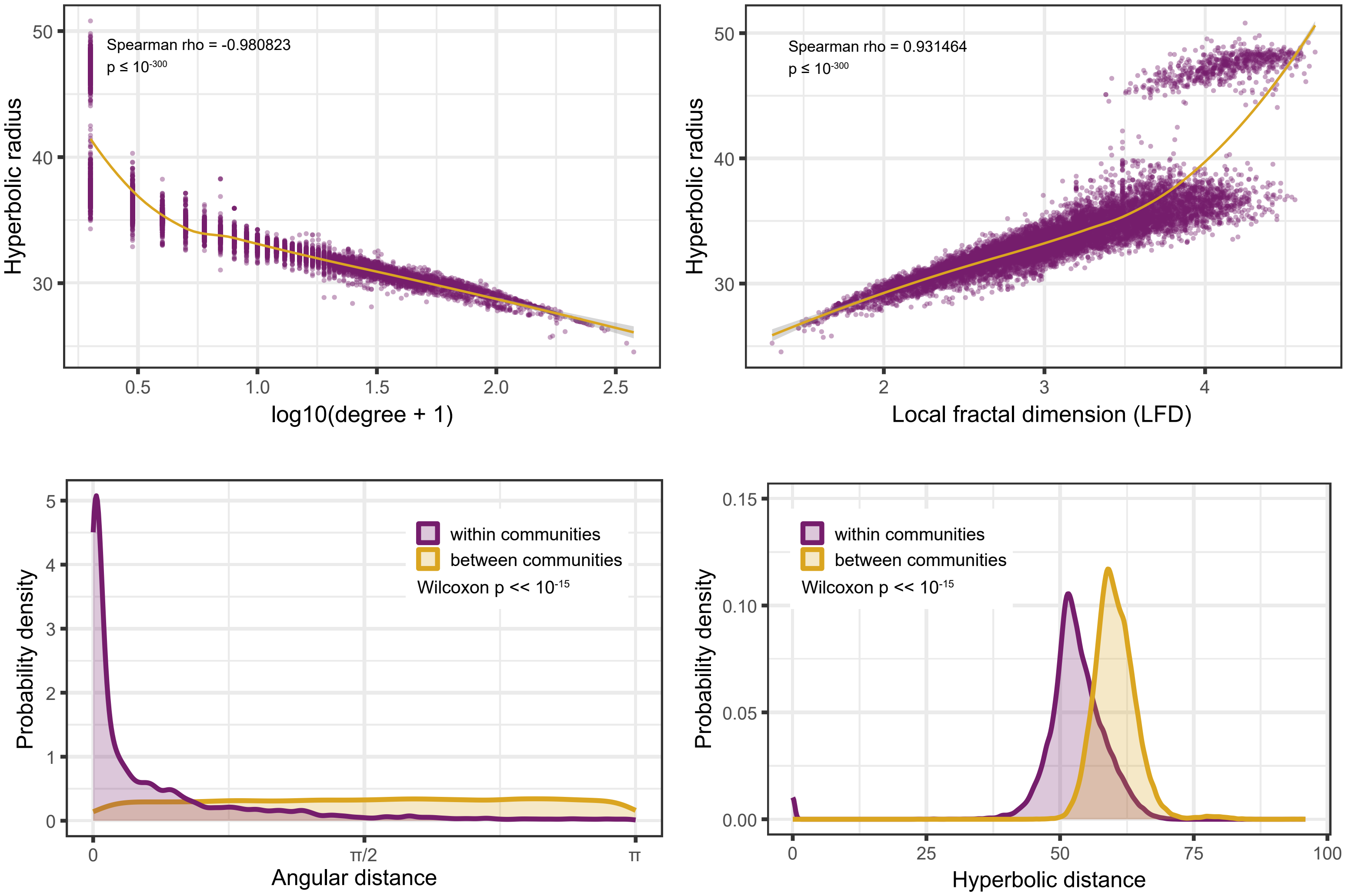

### Supplementary Material

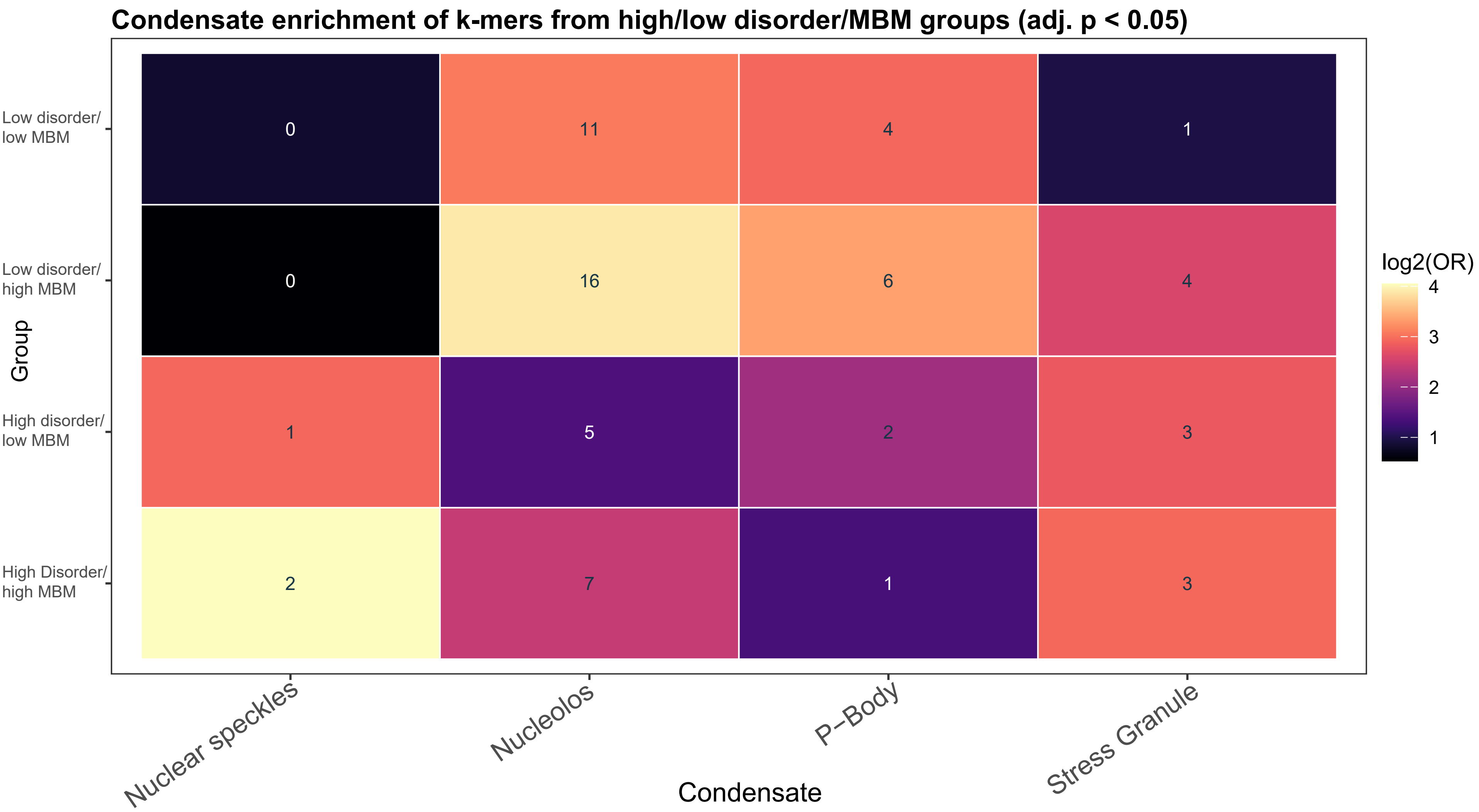
